# The Confessions of a Protein Language Model: Steering, Activations, and What Predicted Confidence Cannot Certify

**DOI:** 10.64898/2026.11.10.765136

**Authors:** Panini Shah, Raj Panchali, Pearl Mody, Mahek Sanghvi, Meera Narvekar, Ruhina Karani

## Abstract

Protein generate-and-screen pipelines use interventions and readouts to select candidates, but rarely test whether these track their intended targets. We audit published repetition steering, ESMFold confidence, and an activation readout. Steering preserves confidence at its operating point, but a fitted cliff follows at *α*_50_ = 1.240× (sweep’s own control: 64.0%). Direction matters at moderate matched-norm push, but not at saturation. Unsteered generator activations predict the confidence gate (ROC-AUC 0.786 ± 0.004), supporting cost-aware, model-specific triage. We also replicate established length sensitivity, through which shortening can manufacture false nulls. Yet on composition- and length-matched design–scramble pairs, no balanced-accuracy interval excludes chance for experimental stability. Predicting the gate does not establish foldability. We should test what a readout measures before using it to select. Code and data: https://github.com/paninishah/plm-steering-audit

## 1 Introduction

Protein design seeks physically meaningful candidates for folding or experimental testing. Pipelines screen language-model proposals with cheap readouts, but rarely test whether steering acts through its target or confidence reflects stability. We audit these assumptions for published repetition steering [20] and mean ESMFold pLDDT, then test confidence and activation readouts against measured protease stability [13].

### Steering window

Our independent readout supports confidence preservation at the mapped operating point, but a fitted dose cliff limits this window.

### Direction specificity

At moderate matched-norm push, the constructed vector causes less collapse, with non-overlapping confidence intervals. Random steering ties it at saturation, masking direction specificity.

### Length sensitivity

Paired truncation raises pLDDT, replicating established length sensitivity [11]. Shortening can therefore create a false null when evaluating an intervention.

### Activation-based triage

Applying established confidence prediction, we predict the ESMFold gate from unsteered activations for model-specific, cost-aware triage.

### Experimental stability

Neither pLDDT nor activations reliably identifies the more stable member of matched design–scramble pairs. Predicting the gate does not establish physical stability.

### Related work

We build on protein steering [6; 20] and non-unique steering directions [17]. Prior work documents pLDDT’s length sensitivity [11] and predicts fold confidence from language-model embeddings [2]. Sparse probing and protein SAE interpretation [15; 9; 19; 10; 14] inform our replication of the language-model sparse-versus-raw result [7]. We apply these methods to audit a published claim (Appendix G).

## 2 Setup

We study ProtGPT2 and cross-check ZymCTRL, RITA-small, p-IgGen, Mistral-Prot-134M, and ProGen2-small. We define collapse as ESMFold’s mean per-residue pLDDT < 60 (appendix results repeat at a secondary threshold of < 70)—confidence, not agreement with experimental structure.

We form a contrastive mean-activation difference *v* from sets matched on foldability utility but separated by repetition. We inject *h* + *αv* at layer 12 and probe layer 30. We measure repetition as the continuation’s distinct-3-mer fraction after removing whitespace.

Doses are multiples of a reference norm (583.998); comparisons spanning layers or models match relative push instead (Section 4).

Four unsteered control arms (53.2%, 62.0%, 64.0%, 52.0%) come from distinct pools, so we compare each intervention only with its own run’s control (Appendix A).

## 3 Repetition does not predict the confidence gate

We first ask whether repetition, the published method’s target, tracks structural confidence. In an *N* = 50 unsteered control set from one generator (ProtGPT2) and one prompt family (random 10–15 residue prefixes cut from real UniProt reference proteins), the association is positive but nonsignificant: distinct-3-mer fraction and confidence have Pearson *r* = +0.2125, *t*(40) = +1.375, and *p* > 0.05. The correlation uses 42 of 50 rows after excluding 8 with empty generated fields.

The two most repetitive sequences have the lowest and highest structural confidence in the control set. Their distinct-3-mer fractions are 0.339 and 0.365, while their pLDDT values are 28.85 and 85.87. The gaps are 0.026 in repetition and 57.03 in pLDDT. The next-lowest fraction is 0.468, clearly separating these two. Figure 2 (left) shows the pair and Figure A5 their folded structures.

**Figure 1:**
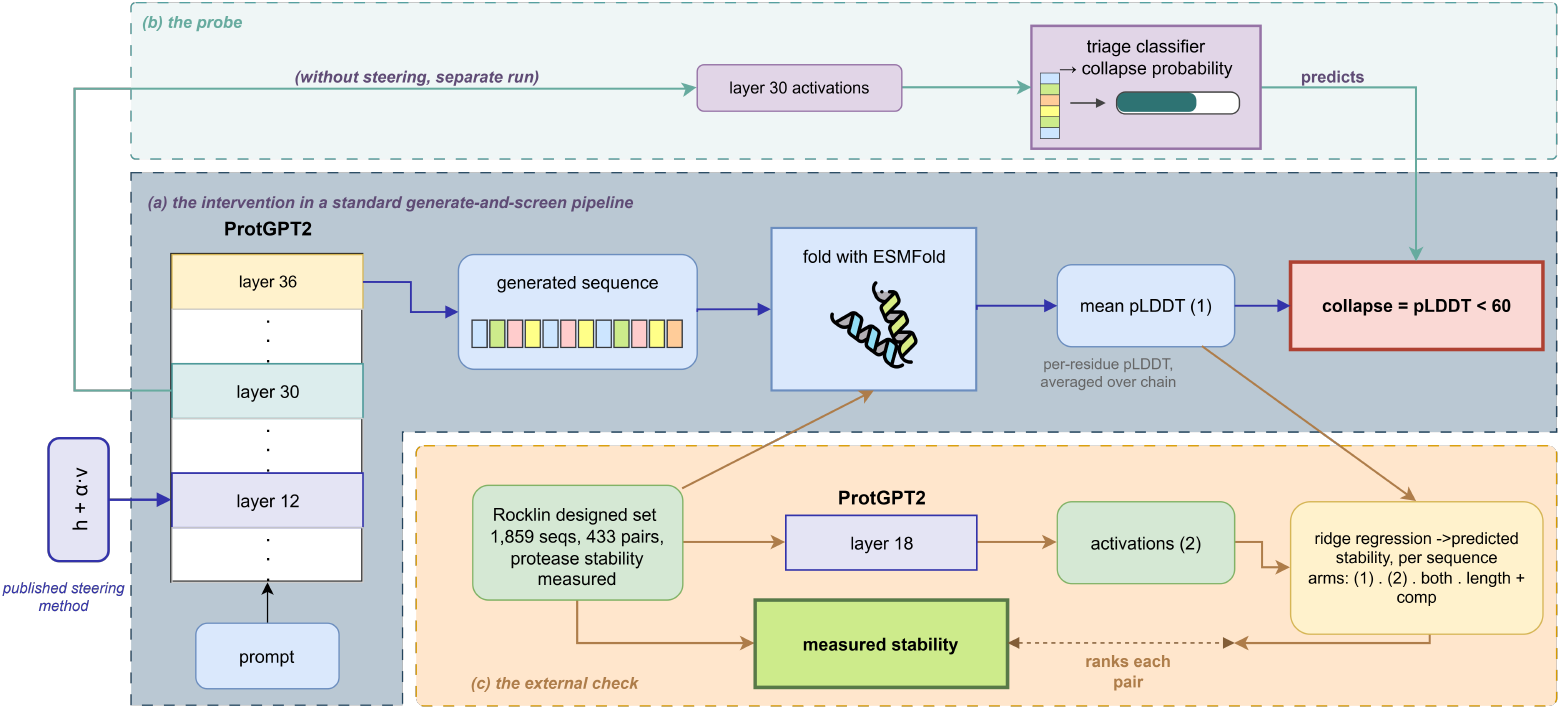
Both arms share one pipeline. (a) ProtGPT2 generates sequences, ESMFold scores them, and mean chain pLDDT defines the confidence gate at pLDDT < 60. The published steering method injects at layer 12 and targets sequence repetition, which Section 3 tests against the gate. (b) On a separate unsteered run, layer-30 activations feed a triage classifier that predicts the gate before folding; it anticipates the structure predictor’s verdict, not physical validity. (c) An external designed set enters at ESMFold and is ranked against measured protease stability by four ridge-regression arms.

**Figure 2:**
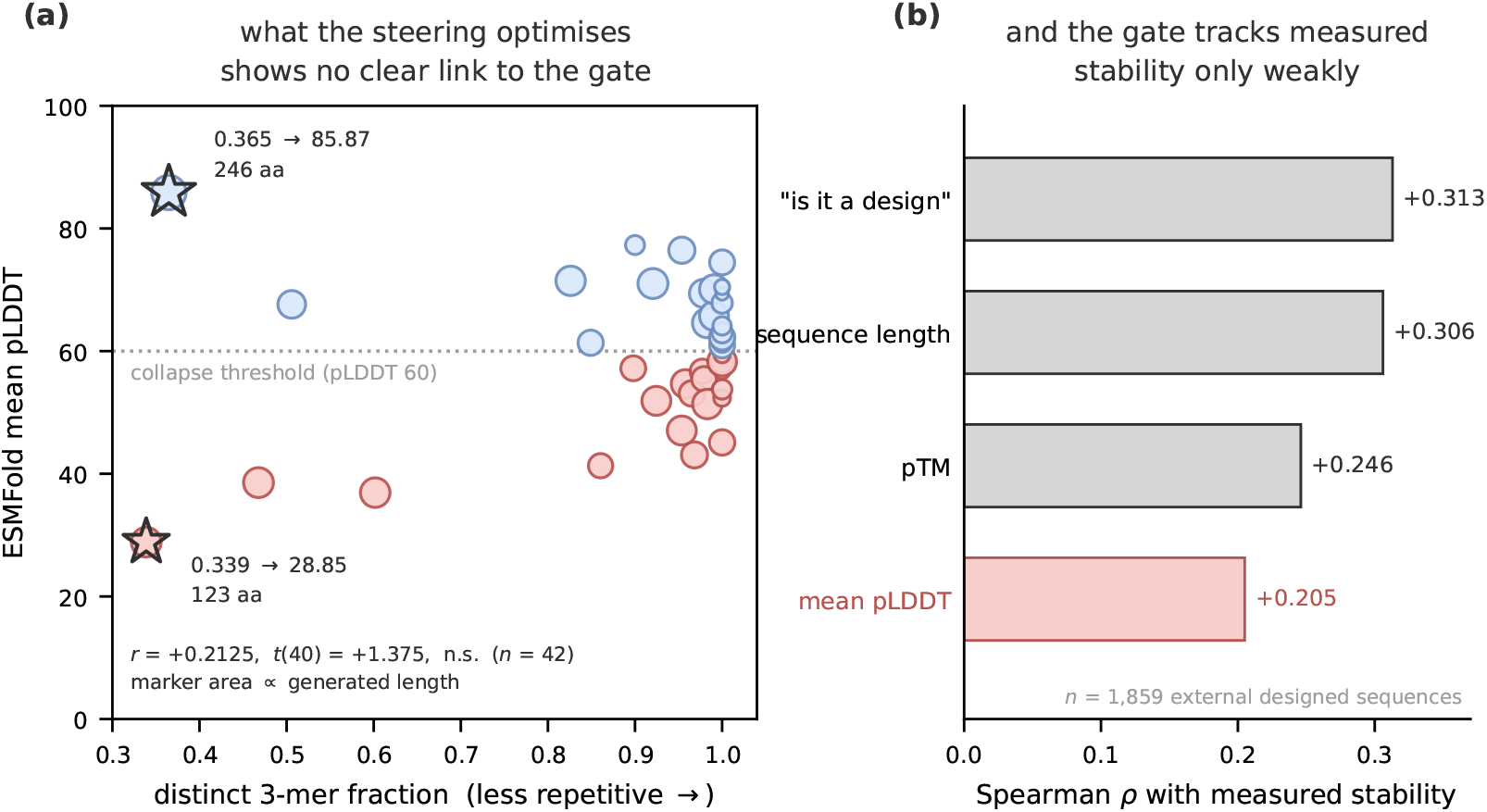
(a) *N* = 50 unsteered ProtGPT2 control set: distinct 3-mer fraction (less repetitive to the right) against ESMFold mean pLDDT. Marker area is proportional to generated length, colour marks the pLDDT 60 gate, and stars mark the two most repetitive sequences, which are the set’s worst- and best-folding members. The correlation uses 42 of the 50 rows, excluding 8 with empty generated fields. (b) 1,859 external designed sequences: Spearman rho between each readout and measured stability. Populations and labels differ between panels, so the two are not directly comparable. The balanced-accuracy intervals discussed in the text come from the matched-pair analysis of Section 7, not from either panel; none excludes chance. Neither result invalidates confidence-gate triage.

Length also affects this readout. A controlled random-sequence sweep on AlphaFold demonstrated pLDDT’s length sensitivity in 2022 [11]. We replicate it in ESMFold by truncating the same 50 sequences to 25, 40, 60, and 80 residues, giving pLDDT values of 61.85, 58.94, 55.91, and 56.65. The paired short-versus-long contrast is 25 versus 80 residues: +5.21 pLDDT (Wilcoxon *p* = 0.0162), with 32 of 50 sequences scoring higher at 25 residues. The difference, the test, and the count all use this pair. Figure A6 plots the pLDDT trend.

The confidence readout therefore also scores an intervention’s effect on generation length. In our steering evaluation, output shortened from 66.8 to 25.5 residues yet appeared unaffected under the confidence readout. Length sensitivity is a plausible explanation for that null, though these data do not establish that shortening concealed a steering effect. Matching length post hoc does not fix this because length mediates the intervention’s effect rather than acting as a nuisance covariate.

## 4 Steering collapses structural confidence above a measured dose threshold

### Dose normalization

We next test how steering affects the gate. Following established activation-norm normalization, we report relative push *α*_rel_ = ‖*v*_inject_ ‖*/* ‖*h*‖ and match it across layers and models, measuring ‖*h*‖ at each injection site on the same 40 real UniProt fragments. At a fixed absolute norm, nominal “1 ×” instead spans 41,033× in relative push. Even at matched push, ZymCTRL needs 2.6–2.8× ProtGPT2’s dose to collapse. Matching makes conditions comparable, not responses equivalent.

### Dose response

For the ProtGPT2 sweep (*N* = 50 per dose), we pin the logistic floor to its own unsteered 64.0% control, giving *α*_50_ = 1.240×, slope *k* = 9.48, and bootstrap 95% CI [1.14, 1.32] for *α*_50_. The largest tested dose not significantly different from control is 1.25× (Fisher *p* = 0.0705); 1.50× reaches 100.0% collapse (*p* < 0.0001). At 0.50× and 0.75× collapse is numerically lower (50.0% each), but the differences are not significant (*p* = 0.2254 each, intervals overlapping control). They reflect noise at *N* = 50, not a protective dip.

**Table A1:**
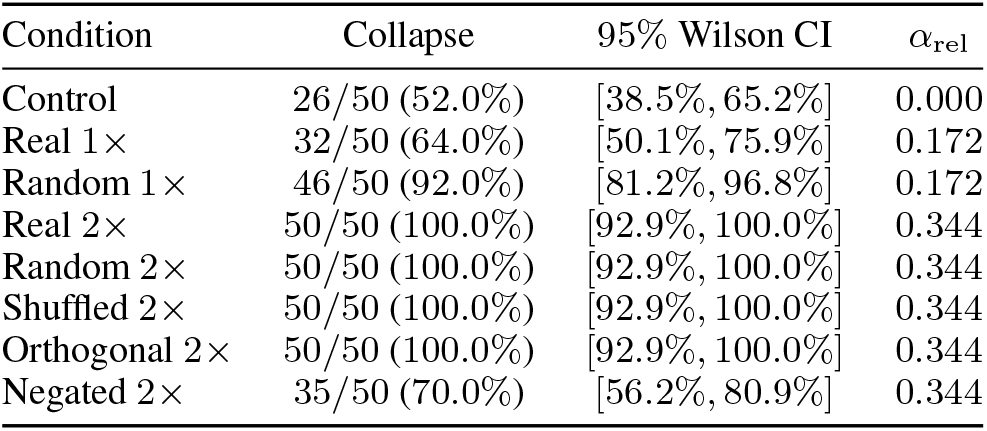
NB35 ProtGPT2 direction controls, *N* = 50 per condition; vector norms match within each push level. This experiment’s own control is 52.0%. Orthogonal 2× is from refolding; at 1 ×, random and real intervals do not overlap.

### Published operating point

Mapping the published operating point below the cliff predicts collapse at the control level, qualitatively confirming preservation. Its stated upper bound straddles the cliff (Table A2). This independently cross-checks rather than refutes the result. The published ablation already acknowledges that repetition control saturates while utility declines beyond approximately *α* = 1–2 [20].

### Direction controls

We test direction specificity in a run with its own 52.0% control. At 2×, random, coordinate-shuffled, and orthogonal directions match the real vector’s 100.0% collapse (Fisher *p* = 1.0000). At matched-norm 1 ×, random gives 92.0% (95% CI [81.2%, 96.8%]), versus real 64.0% ([50.1%, 75.9%]). The non-overlapping intervals indicate least collapse for the real vector at moderate push. Saturation masks this dependence. Negation at 2× gives 70.0%, significantly below forward steering (*p* < 0.0001), but not significantly above its own control (*p* = 0.1004).

### Negative results and reproducibility

Steering toward foldability does not help across two independent attempts; the orthogonalized version is significantly worse than its own control at 1 × (Fisher *p* = 0.0002) and 2 × (*p* < 0.0001). At matched push, 5 of 6 models show no layer effect (Appendix B.5).

## 5 Deep activations predict collapse early enough to triage

Although repetition does not reliably predict the gate, activations may support triage. Prior work predicts AlphaFold2 pLDDT from ESM2 embeddings for 1.5M natural sequences at 0.007 s each [2]. We apply this idea to unsteered ProtGPT2 generations under ESMFold, using binary triage with abstention rather than regression.

### Predictive signal and controls

Deep activations predict the gate with layer-30 ROC-AUC 0.786±0.004 at *N* = 1200, replicating 0.772 ±0.034 at *N* = 400 (means and standard deviations across 20 repeated stratified 70*/*30 splits). Only *N* = 400 supplies matched controls: permuted labels 0.496 ±0.044, entropy 0.586 ±0.040, raw layer 12 0.712 ±0.041, versus layer 30 0.772 ±0.034. Unlike the saturated direction controls, these simpler baselines lose performance. The larger-pool result is a separate replication, not a matched control. Figure A4 plots predicted-collapse precision versus coverage (a) and collapse-prediction AUC from the first *K* residues’ activations (b).

### Selective prediction and early abort

At *N* = 1200, top-decile predicted-collapse precision averages 98.1% across the 10 repeats (sd 1.7 points), with a Wilson 95% CI of [87.9%, 99.6%] computed from the mean call counts of 42 of 43. In the separate early-abort pool (*N* = 250, 15 repeated stratified 70*/*30 splits), the first 30 residues retain 94.9% of full-sequence AUC (0.740 versus 0.780); ZymCTRL replicates retention at 98.1%.

### Sensitivity and transfer limits

The sub-clinical sensitivity test is a null result: the readout does not demonstrate greater sensitivity than the structural endpoint. Zero-shot cross-model transfer ROC-AUC is 0.409, below chance, indicating a model-specific signature under this distribution shift.

## 6 What triage costs and saves

The value of triage depends on cost: a fold costs 77× as much as a classifier call. Batching speeds up generation by 5.80× but length-sorted folding by only 1.33×, raising folding’s share of the pipeline from 81% to 95%. With realistic batching, triage saves roughly 27% of pipeline compute while retaining 81% of candidates that pass the confidence gate.

## 7 What the confidence score does not measure

Useful gate prediction leaves a separate question: do these readouts track experimental stability? We test 433 design–scramble pairs with protease measurements [13], matched on length and composition but differing in amino-acid order. “Always pick the design” has 50.0% balanced accuracy. Our label subtracts a scramble-trained prediction of unfolded-state protease log_10_ EC_50_ from each measurement, correcting length and composition by construction. Identical paired features force the composition-only arm to tie at 50.0%, with CI [50.0%, 50.0%].

ESMFold assigns higher confidence to designs than scrambles: mean pLDDT is 75.5 versus 63.2 for scrambles (+12.3; Mann–Whitney *p* = 5.33 ×10^*−*96^). Across 1,859 sequences, stability correlates with length at Spearman *ρ* = +0.306 versus +0.205 for mean pLDDT; these point estimates alone do not establish a significant difference.

Neither pLDDT nor activations reliably distinguishes the experimentally more stable member of a matched pair. Balanced accuracy (95% CI) is 48.0% for pLDDT ([44.8%, 51.4%]), 51.4% for activations ([48.7%, 54.1%]), and 49.9% for both ([47.3%, 52.5%]).^2^ Raw accuracies (60.3%, 64.7%, 63.5%) exceed 50.0%, but not the always-design rule’s 65.6%. No balanced-accuracy interval excludes chance. The models favor designs, with accuracy 87.3%–94.0% when the design wins versus 6.0%–8.7% when the scramble wins. Adding activations to pLDDT gives no significant improvement (paired Δ = +0.019, CI [*−*0.024, +0.059]).

Gap-stratified point estimates suggest coarse rather than fine stability sensitivity, but no gap’s interval excludes chance (Figure A7).

## 8 Limitations

Our classifier uses thresholded confidence labels rather than measured folding, so it predicts the gate and its errors, not physical validity (Sections 3 and 7). Samples are small: *N* = 50 per steering condition and *N* = 1200 labeled sequences for prediction. The sparse comparison uses a workshop-scale dictionary of 2,560 latents. Reproducibility is moderate (split-half direction cosines 0.74–0.84; worst 0.41), so small effects from single runs remain provisional until replicated. Dose mapping assumes a utility-matched proxy for the unreported norm, magnitude dominance supported only at saturation, and comparability despite unknown injection layer and different instruments. Scope remains limited to one generator and prompt family despite cross-model checks; the matched pair depends on the prompt. Sections 4 and 5 report both arms’ negative results.

## 9 Conclusion

A generate-and-screen pipeline inherits trust: the intervention should act through its target, the readout should reflect physical quality. We tested each link by removing what it depends on. Steering holds only inside a window: below the cliff at *α*_50_ = 1.240× it preserves confidence, but above saturation an arbitrary matched-norm direction does equal damage, turning concept control into magnitude. The readout has an axis of its own, since pLDDT rises under truncation, so an intervention that shortens output can appear inert. Activations are the link that holds, anticipating the gate early enough to abandon a candidate mid-sequence and save roughly 27% of compute at 81% recall. That usefulness stops there: on 433 matched design–scramble pairs, no balanced-accuracy interval excludes chance, and repetition showed no significant association with the gate. Each link is sound for what it measures; the error is inheriting trust across them.

## A Experimental setup and reproducibility

ProtGPT2 [4] is the principal generator. Cross-model checks use ZymCTRL [12], RITA-small [5], p-IgGen [16], Mistral-Prot-134M, and ProGen2-small. ESMFold [8] supplies raw per-residue pLDDT, averaged over the folded chain. The primary collapse label is mean pLDDT < 60; the secondary label is < 70. Here, “foldable” means passing the confidence gate, not experimentally verified folding. The prediction arm runs on *unsteered generations*.

Following established contrastive activation steering [6; 20], we form a mean-difference vector from candidate sets matched on foldability utility and separated by repetition. ProtGPT2 receives *h* + *αv* at layer 12. Repetition is the distinct-3-mer fraction of the generated continuation after whitespace removal. Nominal doses are multiples of `REFERENCE_NORM=` 583.998. Relative push is *α*_rel_ = ‖*v*_inject_‖ */*‖*h*‖ . The cross-model normalization uses forward hooks at each injection site to measure mean per-position activation norms on a common set of 40 real UniProt fragments. Relative push depends on the stated probe set; it is not a model-independent constant or a cross-model predictor of collapse.

The extraction convention `hidden_states[L]` returns the input to block *L*, so vectors built this way sit one block earlier than the injection hook. We measured the difference, which changes no result.

The prediction readout uses layer-30 activations. Its processing pipeline standardizes features, reduces them with PCA-20, and applies a random forest [1]. The full layer sweeps also evaluate support-vector machines (SVMs). The sparse comparison uses matched feature budgets and classifiers, with latent selection fitted inside each cross-validation fold. Its point estimates have no supplied uncertainty estimates and support no significance claim.

Headline and layer-sweep ROC-AUCs are means ± standard deviations across 20 repeated stratified 70*/*30 resampling splits, *not confidence intervals*. Risk–coverage instead uses out-of-fold probabilities from `cross_val_predict`. The *N* = 400 analysis uses 10-fold cross-validation; the NB59 *N* = 1200 risk–coverage analysis uses 10 repeats of stratified 10-fold cross-validation. The early-abort ProtGPT2 pool is *N* = 250, with 15 repeated stratified 70*/*30 splits. These evaluation schemes and pools are not interchangeable.

Collapse-rate intervals and reported precision intervals are 95% Wilson intervals [18]. Steering comparisons use Fisher exact tests with the comparator identified explicitly. For the NB37 dose fit, we fix the logistic floor at that sweep’s own 64.0% control rate and obtain the threshold’s 95% interval by bootstrap resampling. These proportion and dose-fit intervals imply no interval for an AUC standard deviation.

### Control variability

The NB37 dose sweep has its own 64.0% control; the NB35 direction run has its own 52.0% control. The separate NB03 *N* = 50 control is 62.0% with 95% CI [48.2%, 74.1%]. The *N* = 400 unsteered reference is 53.2%. These are distinct pools. We compare each intervention only with its own run’s control and report the spread without pooling or smoothing it.

## B Steering arm: full results

### Reproducibility and cross-model limits

Three models are inert at matched dose. Direction reproducibility is moderate, with split-half cosines of 0.74–0.84 and a worst split of 0.41. Our disjoint-half comparison is stricter than the literature’s subset-versus-reference test, whose reference contains the evaluated subset.

### B.1 Dose response

The NB37 sweep’s own control is 64.0%. Its fitted threshold is *α*_50_ = 1.240× (bootstrap 95% CI [1.14, 1.32]), with slope *k* = 9.48. The largest tested dose not significantly different from control is 1.25× (Fisher *p* = 0.0705); 1.50× has *p* < 0.0001. The lower rates at 0.50 and 0.75× are not significant (*p* = 0.2254 each) and are not evidence of a protective dip.

**Figure A1:**
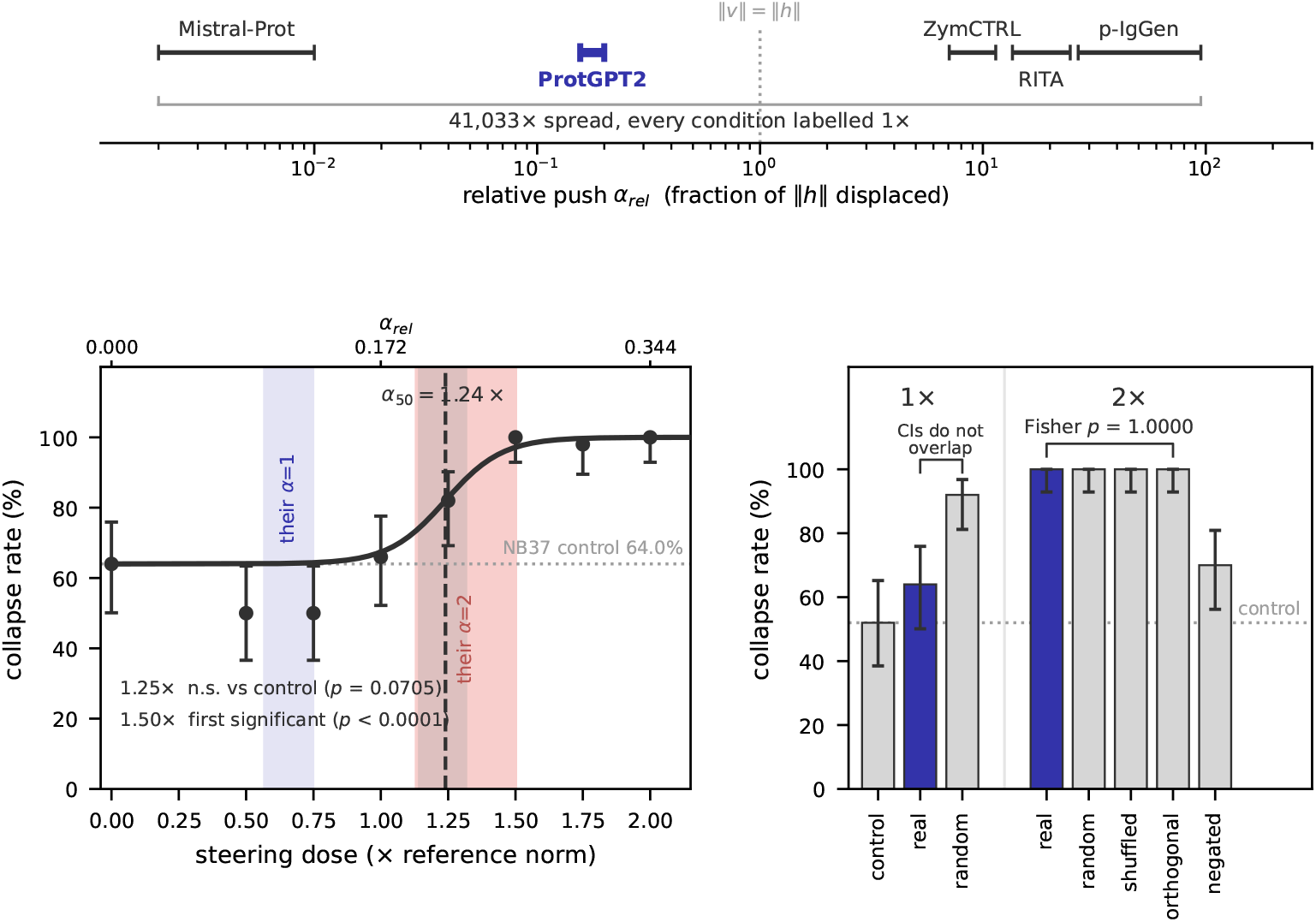
ProtGPT2 (*N* = 50/condition). Top: 1× nominal, 41,033× spread. Left: sweep’s own unsteered control 64.0%; fit *α*_50_ = 1.240×, bootstrap 95% CI [1.14, 1.32]. Bands: published *α* = 1, 2, proxy-mapped, not observed. Right: matched-norm directions; own control 52.0%.

**Table A1:**
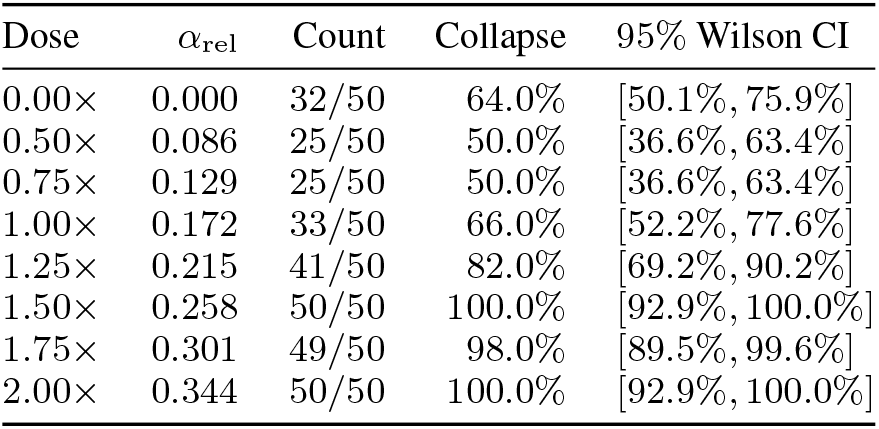
Full NB37 ProtGPT2 layer-12 dose sweep, *N* = 50 per dose (400 folds), evaluated by mean ESMFold pLDDT < 60. Intervals are 95% Wilson intervals. This sweep’s own unsteered control is 64.0%.

| Dose | α<sub>rel</sub> | Count | Collapse | 95% Wilson CI |
|---|---|---|---|---|
| 0.00× | 0.000 | 32/50 | 64.0% | [50.1%, 75.9%] |
| 0.50× | 0.086 | 25/50 | 50.0% | [36.6%, 63.4%] |
| 0.75× | 0.129 | 25/50 | 50.0% | [36.6%, 63.4%] |
| 1.00× | 0.172 | 33/50 | 66.0% | [52.2%, 77.6%] |
| 1.25× | 0.215 | 41/50 | 82.0% | [69.2%, 90.2%] |
| 1.50× | 0.258 | 50/50 | 100.0% | [92.9%, 100.0%] |
| 1.75× | 0.301 | 49/50 | 98.0% | [89.5%, 99.6%] |
| 2.00× | 0.344 | 50/50 | 100.0% | [92.9%, 100.0%] |

### B.2 Published-dose mapping

**Table A2:** NB03 control (*N* = 50): 62.0%, not the dose sweep’s own 64.0%. Cliff row: fitted threshold, no observed collapse rate. Assumptions: utility-matched proxy norm; magnitude dominance only at saturation; comparability despite unknown injection layer and different structure predictors.

| Condition | Dose on our axis | Observed collapse | Mapped prediction |
|---|---|---|---|
| Unsteered control | – | 62.0% | – |
| Published operating point, *α* = 1 | 0.56–0.75× | – | ∼64% |
| Fitted cliff, *α*<sub>50</sub> | 1.240× (CI [1.14, 1.32]) | – | – |
| Published upper bound, *α* = 2 | 1.13–1.51× | – | 73–97% |

### B.3 Direction controls

The NB35 direction experiment has its own 52.0% control. At matched 1× norm, random and real directions differ: 92.0% [81.2%, 96.8%] versus 64.0% [50.1%, 75.9%]. At 2×, real, random, shuffled, and orthogonal directions all give 100.0% collapse (Fisher *p* = 1.0000 for each comparison with real). Negation gives 70.0%: below real (*p* < 0.0001), but not significantly above this run’s control (*p* = 0.1004). The orthogonal condition uses the NB39 refolded result.

**Table A3:**
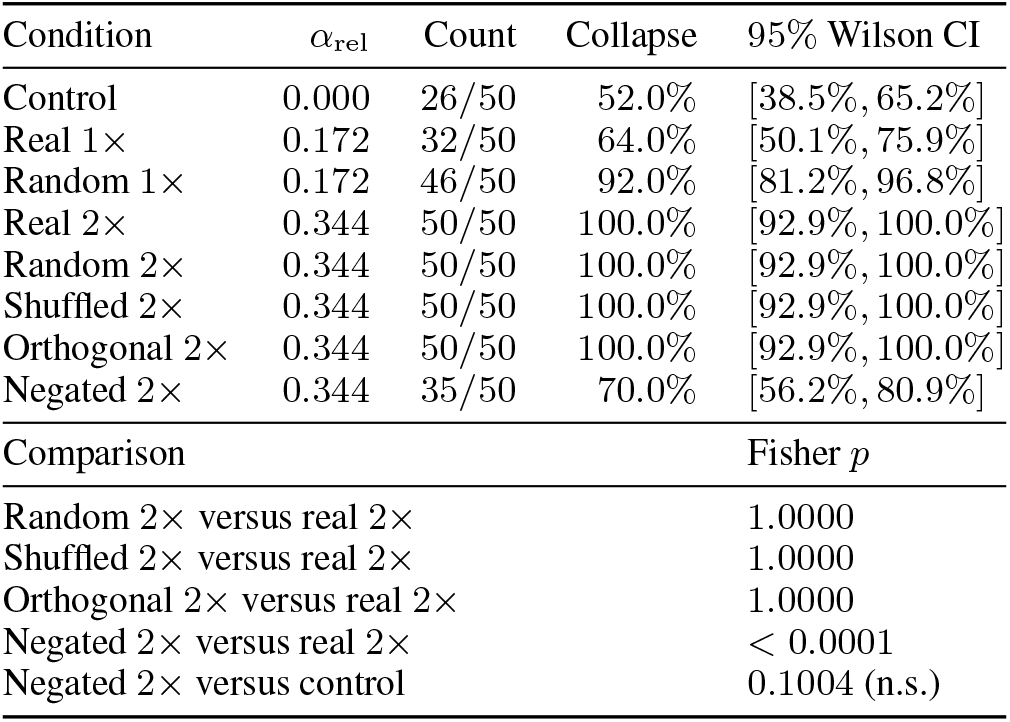
Full NB35 ProtGPT2 layer-12 direction-control pool, *N* = 50 per condition, with the orthogonal condition refolded in NB39. Evaluation uses mean ESMFold pLDDT < 60, 95% Wilson intervals, and Fisher exact comparisons below. Norms are matched within each dose. This run’s own control is 52.0%.

| Condition | α<sub>rel</sub> | Count | Collapse | 95% Wilson CI |
|---|---|---|---|---|
| Control | 0.000 | 26/50 | 52.0% | [38.5%, 65.2%] |
| Real 1× | 0.172 | 32/50 | 64.0% | [50.1%, 75.9%] |
| Random 1× | 0.172 | 46/50 | 92.0% | [81.2%, 96.8%] |
| Real 2× | 0.344 | 50/50 | 100.0% | [92.9%, 100.0%] |
| Random 2× | 0.344 | 50/50 | 100.0% | [92.9%, 100.0%] |
| Shuffled 2× | 0.344 | 50/50 | 100.0% | [92.9%, 100.0%] |
| Orthogonal 2× | 0.344 | 50/50 | 100.0% | [92.9%, 100.0%] |
| Negated 2× | 0.344 | 35/50 | 70.0% | [56.2%, 80.9%] |
| Comparison |  |  |  | Fisher *p* |
| Random 2× versus real 2× |  |  |  | 1.0000 |
| Shuffled 2× versus real 2× |  |  |  | 1.0000 |
| Orthogonal 2× versus real 2× |  |  |  | 1.0000 |
| Negated 2× versus real 2× |  |  |  | < 0.0001 |
| Negated 2× versus control |  |  |  | 0.1004 (n.s.) |

### B.4 Steering toward foldability: both attempts

#### Attempt 1: co-linear

The NB36 foldability vector has cosine +0.9819 with the repetition vector. This nearly re-measures the same direction rather than testing an independent one. Its own control is 52.0%. Differences at the lower doses are not significant; at 2 ×, collapse is 90.0% (Fisher *p* < 0.0001 versus control).

**Table A4:** Attempt 1, NB36 ProtGPT2 layer-12 pool, *N* = 50 per condition. Collapse is mean ESMFold pLDDT < 60; intervals are 95% Wilson intervals and Fisher tests compare each dose with this run’s own 52.0% control. The direction cosine is +0.9819; n.s. denotes nonsignificance.

| Dose | *α*<sub>rel</sub> | Collapse count (rate) | 95% Wilson CI | Fisher *p* |
|---|---|---|---|---|
| 0.25× | 0.050 | 23/50 (46.0%) | [33.0%, 59.6%] | 0.689 (n.s.) |
| 0.5× | 0.101 | 23/50 (46.0%) | [33.0%, 59.6%] | 0.689 (n.s.) |
| 1× | 0.202 | 31/50 (62.0%) | [48.2%, 74.1%] | 0.419 (n.s.) |
| 2× | 0.404 | 45/50 (90.0%) | [78.6%, 95.7%] | < 0.0001 |

#### Attempt 2: orthogonalized

The NB39 follow-up projects out the shared component, leaving cosine −0.0000, orthogonal to machine precision. We rebuild the vector from a fresh 200-candidate pool with 57.0% natural collapse. The intervention run has its own 52.0% control. Rather than a null result, this attempt significantly worsens collapse relative to control at 1× (*p* = 0.0002) and 2× (*p* < 0.0001). Across increasing doses it rises monotonically, 60.0% →70.0% →88.0% → 100.0%. The lower-dose differences are not significant.

**Table A5:** Attempt 2, NB39 ProtGPT2 layer-12 pool, *N* = 50 per condition, using the orthogonalized vector from the fresh 200-candidate pool. Evaluation uses mean ESMFold pLDDT < 60, 95% Wilson intervals, and Fisher tests against this run’s own 52.0% control. The direction cosine is −0.0000; n.s. denotes nonsignificance.

| Dose | Collapse count (rate) | 95% Wilson CI | Fisher *p* |
|---|---|---|---|
| 0.25× | 30/50 (60.0%) | [46.2%, 72.4%] | 0.546 (n.s.) |
| 0.5× | 35/50 (70.0%) | [56.2%, 80.9%] | 0.100 (n.s.) |
| 1× | 44/50 (88.0%) | [76.2%, 94.4%] | 0.0002 |
| 2× | 50/50 (100.0%) | [92.9%, 100.0%] | < 0.0001 |

### B.5 Within-model layer dose asymmetry

**Figure A2:**
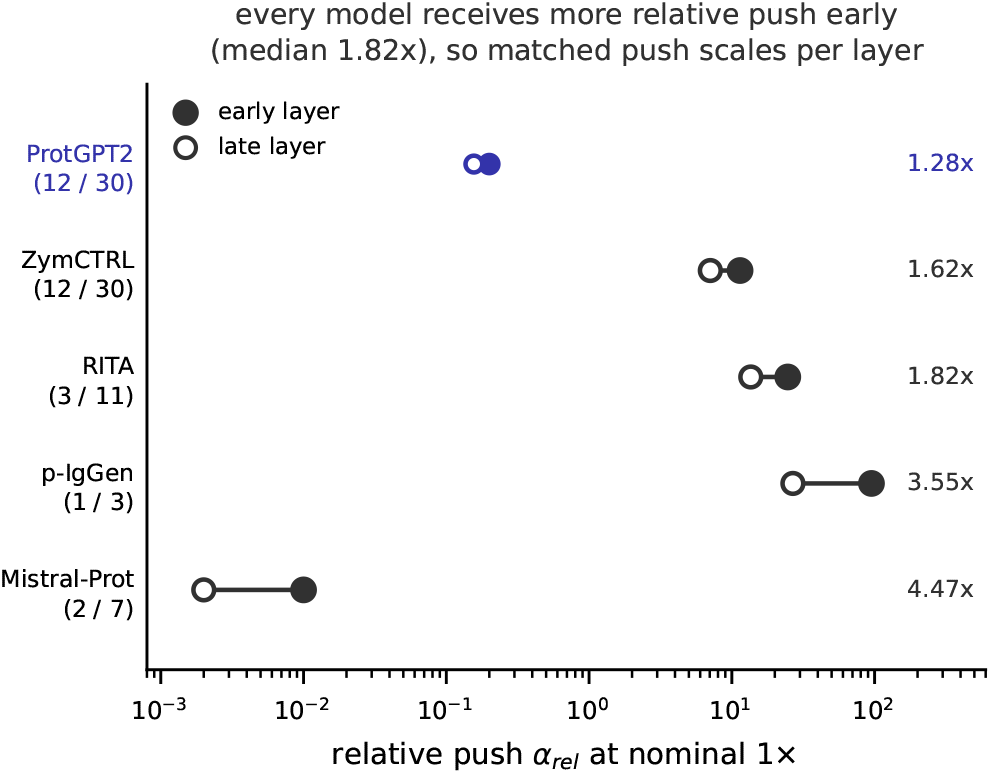
Relative push at early and late injection layers at nominal 1×, measured on the common probe set. Every model’s early layer receives more relative push, by a median of 1.82× ; matched-push testing therefore scales the dose per model and layer.

### B.6 Cross-model matched push

Nominal doses are multiples of a single reference norm, so the same label means a different relative push in every model. Comparisons that span layers or models therefore rescale the injected vector to a common *α*_rel_ measured at each site.

**Table A6:** Relative push at nominal 1×, measured with forward hooks at each injection site on a common set of 40 real UniProt fragments. Every condition in this table is labelled “1×”; they span 41,033×.

| Model | Layer | ‖*h*‖ | raw ‖*v*<sub>*L*</sub>‖ | *α*<sub>rel</sub> at 1× | at 2× |
|---|---|---|---|---|---|
| Mistral-Prot | 7 | 252,032.61 | 5280.0 | 0.002 | 0.005 |
| Mistral-Prot | 2 | 56,439.23 | 3664.0 | 0.010 | 0.021 |
| ProtGPT2 | 30 | 3751.99 | 781.18 | 0.156 | 0.311 |
| ProtGPT2 | 12 | 2921.53 | 439.62 | 0.200 | 0.400 |
| ZymCTRL | 30 | 82.66 | 5.944 | 7.065 | 14.130 |
| ZymCTRL | 12 | 51.14 | 3.357 | 11.419 | 22.839 |
| RITA | 11 | 43.03 | 5.351 | 13.572 | 27.143 |
| RITA | 3 | 23.69 | 2.979 | 24.652 | 49.304 |
| p-IgGen | 3 | 21.81 | 1.832 | 26.774 | 53.547 |
| p-IgGen | 1 | 6.14 | 0.567 | 95.079 | 190.159 |

At p-IgGen layer 1, “1×” adds a vector 95 times larger than the hidden state it is added to; at Mistral-Prot layer 7 it displaces that state by 0.2%. Relative push is not itself a universal dose. ProtGPT2 reaches 90% collapse at *α*_rel_ = 0.400 while ZymCTRL requires 22.8, and LayerNorm downstream partially rescales a large added vector. Relative push allows comparison but does not predict collapse across models. We fit no cross-model dose–response curve because the tested points cluster at 0.16–0.40 and 7–190, with nothing between.

**Table A7:**
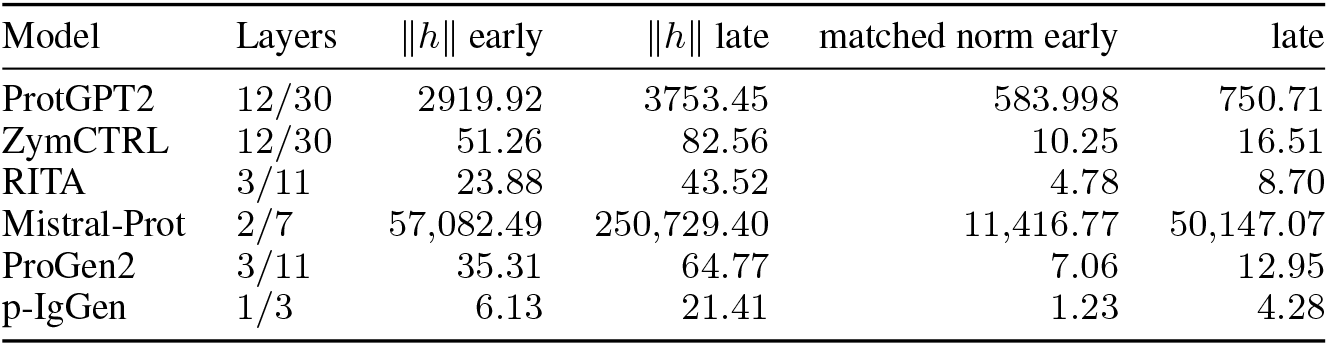
Matched-push calibration. We recompute each layer’s injected norm from its own ‖*h*‖ to hold *α*_rel_ = 0.200, the value ProtGPT2 layer 12 receives at nominal 1× . Norms differ by four orders of magnitude across models. Remeasuring norms within each repair run gives slight differences from Table A6’s probe-set values. Relative push is a measured quantity, not a constant.

| Model | Layers | ‖*h*‖ early | ‖*h*‖ late | matched norm early | late |
|---|---|---|---|---|---|
| ProtGPT2 | 12/30 | 2919.92 | 3753.45 | 583.998 | 750.71 |
| ZymCTRL | 12/30 | 51.26 | 82.56 | 10.25 | 16.51 |
| RITA | 3/11 | 23.88 | 43.52 | 4.78 | 8.70 |
| Mistral-Prot | 2/7 | 57,082.49 | 250,729.40 | 11,416.77 | 50,147.07 |
| ProGen2 | 3/11 | 35.31 | 64.77 | 7.06 | 12.95 |
| p-IgGen | 1/3 | 6.13 | 21.41 | 1.23 | 4.28 |

**Table A8:** ZymCTRL dose ladder at matched relative push, *N* = 50 per condition, with this run’s own 62.0% control. Fisher exact tests against that control. The first significant dose is 3× the matched scale; against ProtGPT2’s own *α*_50_ = 0.2131 this is 2.6–2.8× the relative push.

| Dose | α<sub>rel</sub> | Collapse | Fisher *p* vs control |
|---|---|---|---|
| Control | 0.000 | 31/50 (62.0%) | – |
| 1× | 0.200 | 34/50 (68.0%) | 0.6753 (n.s.) |
| 2× | 0.400 | 37/50 (74.0%) | 0.2837 (n.s.) |
| 3× | 0.600 | 43/50 (86.0%) | **0.0113** |
| 4× | 0.800 | 45/50 (90.0%) | **0.0019** |

Both models collapse under sufficient relative perturbation, but require different amounts. This measured cross-model difference does not demonstrate that ZymCTRL is robust.

## C Prediction arm: full results

Both layer sweeps use the NB08 *N* = 400 unsteered ProtGPT2 pool. The primary threshold has 53.2% collapse; the secondary threshold has 81.0%. These tables compare raw activations and entropy features with random forests and SVMs.

### C.1 Primary threshold

**Table A9:** Internally matched prediction baselines on the same *N* = 400 ProtGPT2 pool, predicting mean ESMFold pLDDT < 60. ROC-AUC values are means ±standard deviations across 20 repeated stratified 70*/*30 resampling splits. The deep-layer reference is 0.772, the smaller-pool replication value, not the separate headline, risk–coverage, early-abort, or compute-pool AUC. The permutation control shuffles labels for the deep-layer readout.

| Readout | ROC-AUC |
|---|---|
| Permutation control | 0.496 ± 0.044 |
| Entropy baseline | 0.586 ± 0.040 |
| Raw activations, layer 12 | 0.712 ± 0.041 |
| Raw activations, layer 30 | 0.772 ± 0.034 |

**Table A10:** Primary-threshold layer sweep, mean ESMFold pLDDT < 60, on the NB08 *N* = 400 unsteered ProtGPT2 pool (53.2% collapse). Random-forest and SVM ROC-AUCs are means ±standard deviations across 20 repeated stratified 70*/*30 splits, not confidence intervals.

| Feature set | Random forest | SVM |
|---|---|---|
| Entropy baseline | 0.586 ± 0.040 | 0.581 ± 0.068 |
| Raw layer 6 | 0.711 ± 0.034 | 0.699 ± 0.047 |
| Raw layer 12 | 0.712 ± 0.041 | 0.709 ± 0.051 |
| Raw layer 18 | 0.705 ± 0.051 | 0.719 ± 0.046 |
| Raw layer 24 | 0.767 ± 0.043 | 0.760 ± 0.049 |
| Raw layer 30 | 0.772 ± 0.034 | 0.757 ± 0.037 |

### C.2 Secondary threshold: not the headline

**Table A11:** At the secondary threshold, mean ESMFold pLDDT < 70, the layer-30 SVM gives 0.786 ±0.035 on the NB08 *N* = 400 unsteered ProtGPT2 pool (81.0% collapse); this is not the headline value. The headline 0.786 ±0.004 instead uses a random forest, pLDDT < 60, and the NB59 *N* = 1200 pool. Both schemes report means ±standard deviations across 20 repeated stratified 70*/*30 splits, not confidence intervals; threshold, classifier, pool, and standard deviation differ.

| Feature set | Random forest | SVM |
|---|---|---|
| Entropy baseline | 0.631 ± 0.043 | 0.632 ± 0.046 |
| Raw layer 6 | 0.731 ± 0.049 | 0.737 ± 0.050 |
| Raw layer 12 | 0.720 ± 0.044 | 0.738 ± 0.044 |
| Raw layer 18 | 0.723 ± 0.051 | 0.755 ± 0.043 |
| Raw layer 24 | 0.763 ± 0.044 | 0.774 ± 0.037 |
| Raw layer 30 | 0.769 ± 0.036 | 0.786 ± 0.035 |

**Figure A3:**
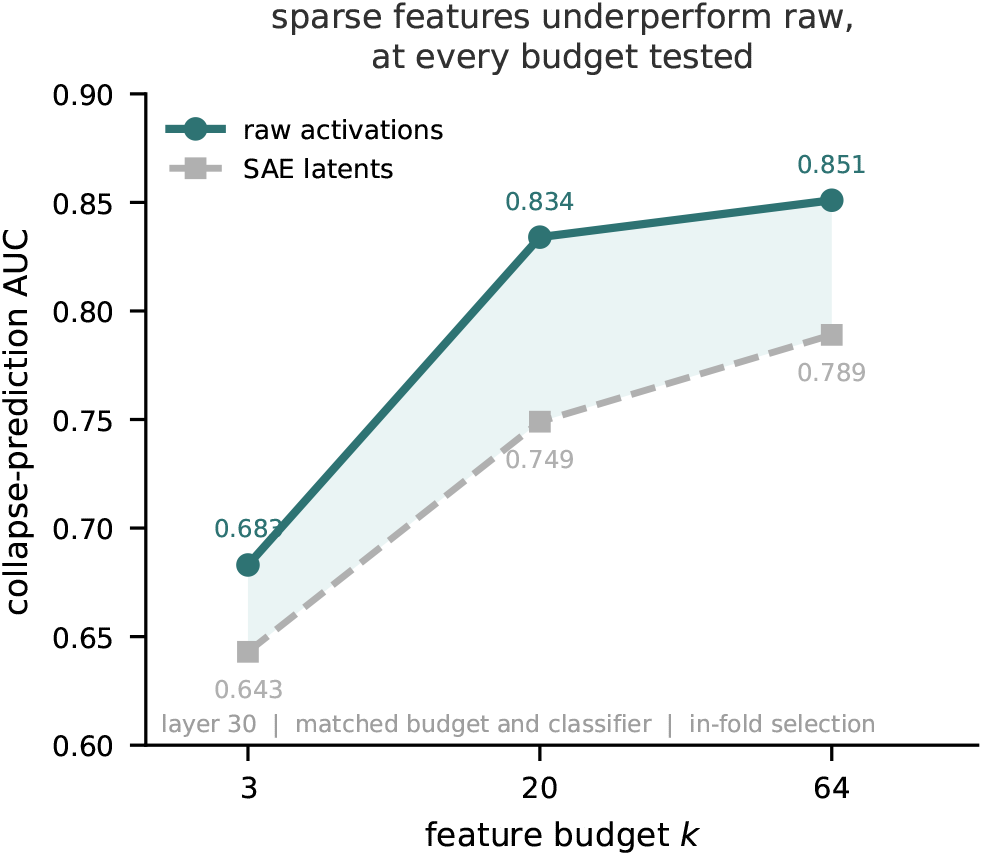
Protein-domain replication of the language-model sparse-probe result. We match layer 30, feature budgets, and classifiers, and fit latent selection within each cross-validation fold. ROC-AUC point estimates lack uncertainty estimates and establish no significance. Dictionary improvement halved the gap from −0.117 to− 0.049, making it an upper bound on sparse-feature disadvantage, not a floor.

### C.3 Matched-budget sparse-feature replication

This protein-domain comparison replicates the sparse-probe disadvantage reported in language models [7]. We match classifiers and budgets and fit latent selection inside each cross-validation fold. The point estimates have no uncertainty estimates and do not establish significance (Figure A3).

Dictionary variance unexplained is 0.093, mean *L*_0_ = 32.0, and 46.7% of 2,560 latents are dead; training uses 721,000 tokens from 12,000 real UniProt proteins. Raw AUC 0.851 is a modeling-choice upper bound, not the headline.

### C.4 Selective prediction and early abort

The left panel uses the NB59 *N* = 1200 unsteered ProtGPT2 pool, whose base rate is 52.2%, with out-of-fold risk–coverage evaluation. The right panel uses the *N* = 250 ProtGPT2 early-abort pool and 15 repeated stratified 70*/*30 splits. Its shaded band is the min–max range across repeats for ProtGPT2 only: per-repeat ranges are unavailable for ZymCTRL and RITA-small. RITA-small stays at chance, so its retention is uninformative. These are model-specific confidence-prediction checks, not experimental stability tests.

**Figure A4:**
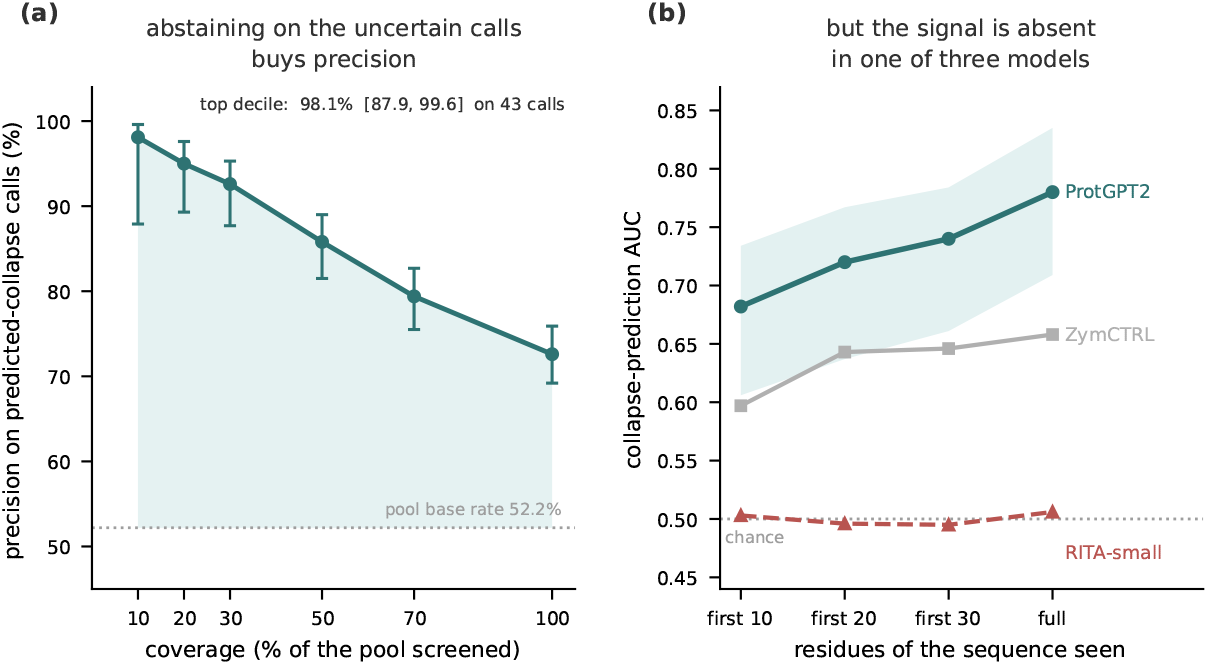
ESMFold confidence on unsteered generations, not experimental stability; base rate 52.2% in (a). In (b), shading is ProtGPT2’s 15-repeat min–max only: ZymCTRL/RITA-small lack per-repeat ranges. RITA-small stays at chance; retention is uninformative. Risk-coverage precisions are means across the 10 repeats; the call counts and their Wilson intervals are computed from the per-repeat counts averaged and rounded, so a precision and its count ratio need not agree exactly.

## D Deployment and compute

Generation supplies activations without another model pass, unlike representation extraction for finished proteins. Each generated candidate needs a fold to obtain a confidence label unavailable from databases, so labeling cost rather than candidate availability limits sample size. Batching leaves pLDDT unchanged.

### D.1 Unbatched cost

The NB45 timing pool contains *N* = 200 generated sequences, of which 198 were folded, with 50.0% natural collapse. Timings use CUDA synchronization on a Tesla T4 after discarding warm-up. ESMFold costs 3.54× a generation and 77× a classifier call. Fold time correlates with length at *r* = +0.892 (mean 58 residues, range 10–210), so its single mean is a simplification rather than a length-independent cost.

**Table A12:** Measured unbatched cost on the NB45 ProtGPT2 pool: *N* = 200 generated, 198 folded. Evaluation uses CUDA-synchronized Tesla T4 timing with warm-up discarded. Fold time has length correlation *r* = +0.892 (mean 58 residues, range 10–210); the precise mean below simplifies length-dependent cost.

| Component | Time | Share |
|---|---|---|
| Full generation (∼ 50 residues) | 0.592 s | 22% |
| ESMFold, one sequence | 2.095 s | 78% |
| Classifier at 30 residues | 0.027 s | ∼ 1% |
| Baseline total | 2.687 s | 100% |

### D.2 Batched throughput

**Table A13:**
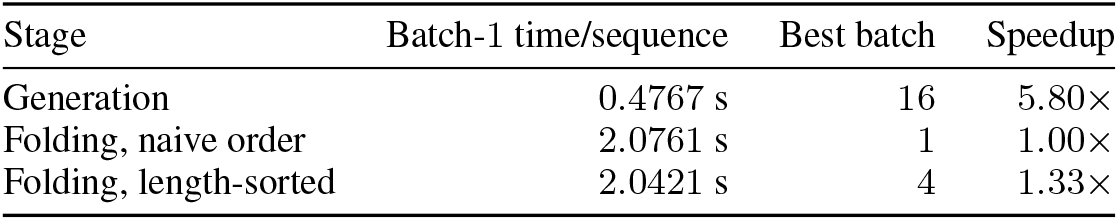
Measured batching benchmark on the fixed *N* = 64 sequence pool, using Tesla T4 generation and ESMFold timing across batch sizes and folding orders. The time column is the batch-1 reference, not the best-batch time. Speedups are measured point estimates without uncertainty intervals.

The same *N* = 64 pool has 0*/*64 label flips under batching and pLDDT correlation 1.0000. Naive folding gives speedups of 0.93× at batch 2 and 0.76× at batch 4. Length sorting is therefore part of the reported batching configuration, not an interchangeable implementation detail. ESMFold runs out of memory at batch 8 on the 16 GB T4; peak memory is 14.47 GB at batch 1, and the maximum feasible batch is 4.

### D.3 Triage policies and batching regimes

The compute evaluation uses the *N* = 198 pool and a single ranked list of pooled out-of-fold scores, with ROC-AUC 0.724. The unbatched early-abort policy costs and batching-regime savings below are separate accounting summaries; their unbatched entries are not interchangeable. Each savings entry retains its corresponding recall of foldable candidates. Charging surviving continuations at unbatched efficiency gives a worst-case saving of 21.1% at 81% recall of foldable candidates.

**Table A14:** Unbatched early-abort policies at 30 residues on the NB45 *N* = 198 folded compute pool. Evaluation ranks pooled out-of-fold scores (ROC-AUC 0.724) and accounts for cost per generated candidate. Each row pairs compute saved with recall of candidates passing the confidence gate.

| Fold budget | Foldable recall | Cost/candidate | Compute saved |
|---|---|---|---|
| 30% | 46.5% | 1.098 s | 59.1% |
| 50% | 69.7% | 1.560 s | 41.9% |
| 70% | 80.8% | 2.021 s | 24.8% |

**Table A15:**
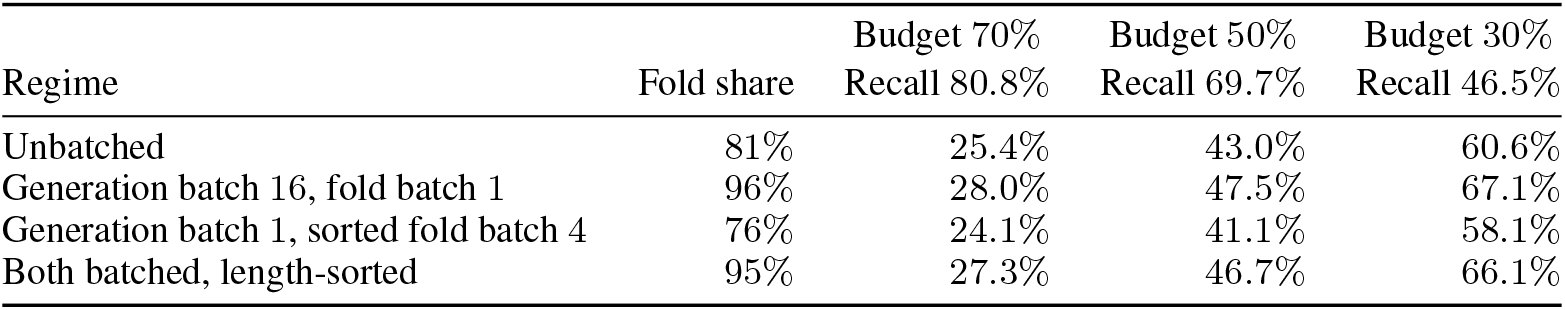
Batching-regime savings for triage on the *N* = 198 compute pool, evaluated from a pooled out-of-fold ranking (ROC-AUC 0.724) and the fixed *N* = 64 batching benchmark. Each savings column identifies its fold budget and recall of foldable candidates. These regime projections are separate from the unbatched early-abort costs in Table A14.

|  |  | Budget 70% | Budget 50% | Budget 30% |
|---|---|---|---|---|
| Regime | Fold share | Recall 80.8% | Recall 69.7% | Recall 46.5% |
| Unbatched | 81% | 25.4% | 43.0% | 60.6% |
| Generation batch 16, fold batch 1 | 96% | 28.0% | 47.5% | 67.1% |
| Generation batch 1, sorted fold batch 4 | 76% | 24.1% | 41.1% | 58.1% |
| Both batched, length-sorted | 95% | 27.3% | 46.7% | 66.1% |

## E The repetition counterexample

The structural pair comes from the unsteered *N* = 50 control set. The repetition measure uses generated continuations, whereas ESMFold receives full chains including the prompt. Similar repetition does not certify similar structural confidence.

**Figure A5:**
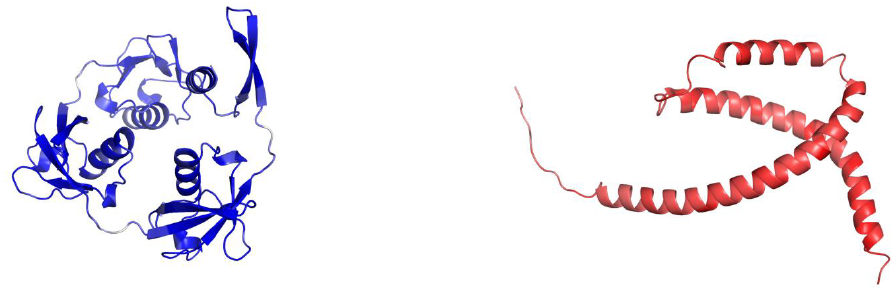
Both panels use one shared zoom scale and one shared per-residue pLDDT colour scale (red = low confidence, blue = high). **Left:** the confident 258-residue chain, mean pLDDT 85.87. **Right:** the collapsed 135-residue chain, 28.85 — shorter, yet the physically larger object (Table A16), which is why independent panel fitting would destroy the contrast. Both are folded chains including the prompt; repetition is measured on their 246- and 123-residue generated continuations. These are predicted structures, not experimental measurements.

**Table A16:**
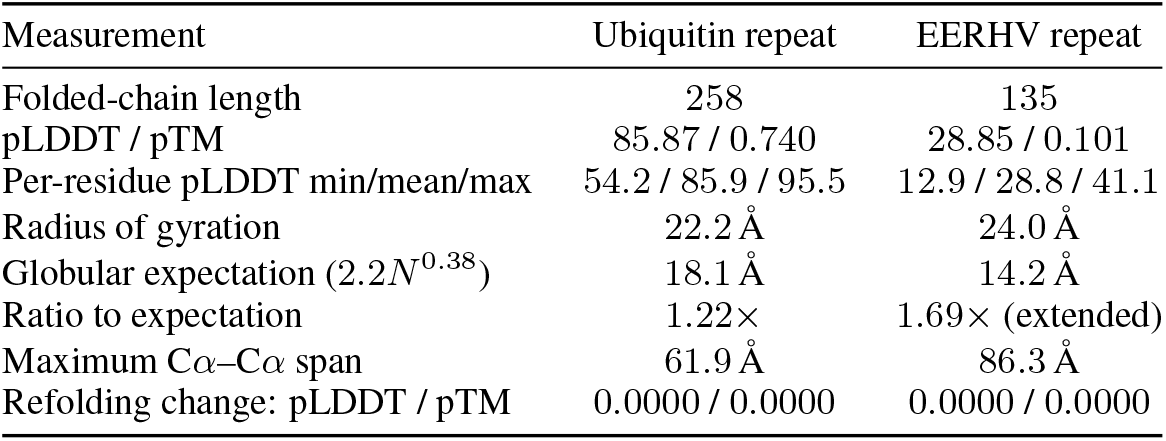
Geometry of the matched pair from the unsteered *N* = 50 ProtGPT2 control pool. Evaluation uses ESMFold scores, folded-coordinate geometry, and deterministic refolding. The reported 85.87 pLDDT is the all-atom per-residue mean, not the C*α*-only mean. Both structures refold bit-for-bit: changes in pLDDT and pTM are 0.0000.

### E.1 Paired truncation

## F Experimental stability evaluation

This evaluation uses experimental measurements from [13], not confidence labels. The score subtracts predicted unfolded-state protease log_10_ EC_50_ from the measured value, using an unfolded-state model parameterized on scrambled sequences. The matched design–scramble pairs have identical length and composition. The composition-only arm therefore ties at 50.0%, with CI [50.0%, 50.0%]. No balanced-accuracy interval excludes chance.

**Figure A6:**
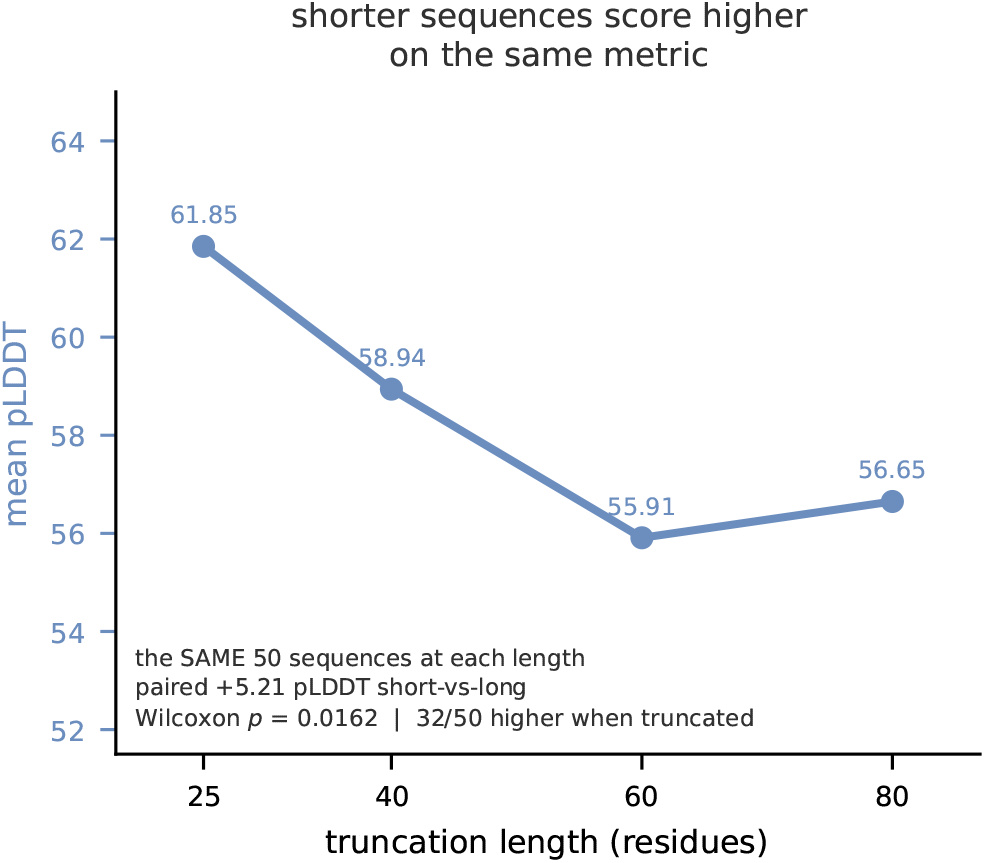
We fold the same 50 sequences at four truncation lengths. Shorter sequences score higher, with paired +5.21 pLDDT short-versus-long, Wilcoxon *p* = 0.0162, with 32*/*50 scoring higher when truncated. We replicate the established AlphaFold result with a paired within-sequence design and examine its consequence for intervention evaluation.

### F.1 Gap-stratified discrimination

**Figure A7:**
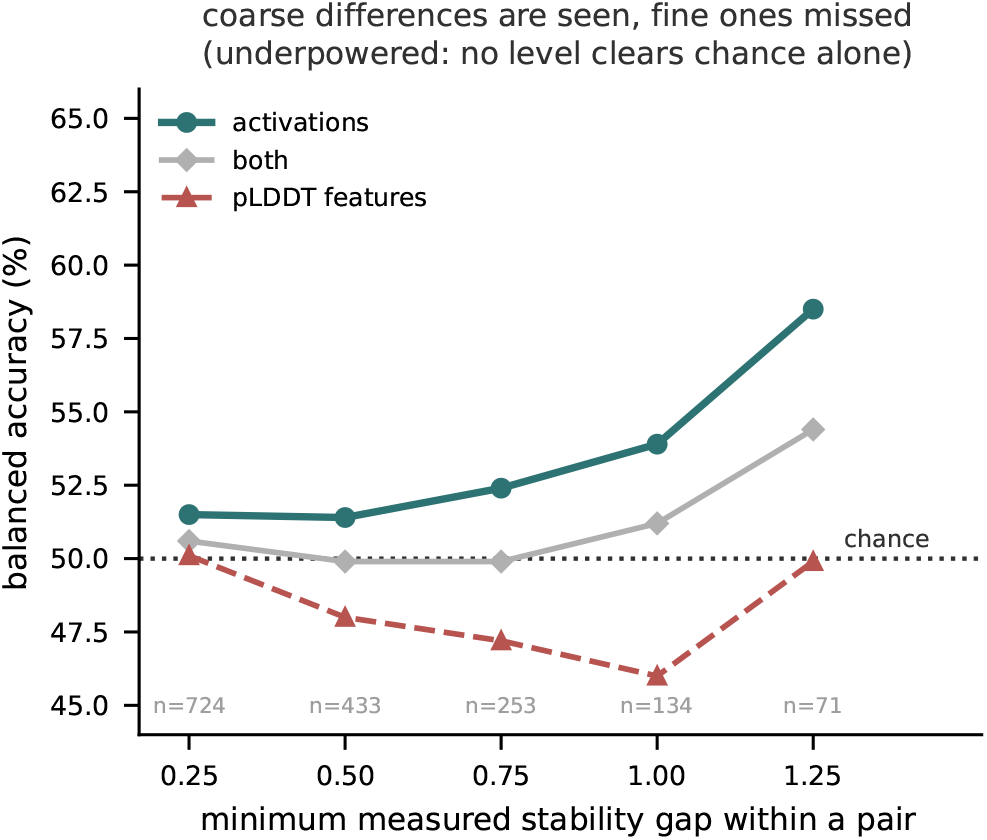
Balanced accuracy against the minimum measured stability gap within a pair. Point estimates suggest sensitivity to coarse rather than fine stability differences, but no gap level independently excludes chance. The pair count falls from 724 to 71 across the sweep. This does not establish a reliable increase.

### F.2 Matched-pair results

**Table A17:**
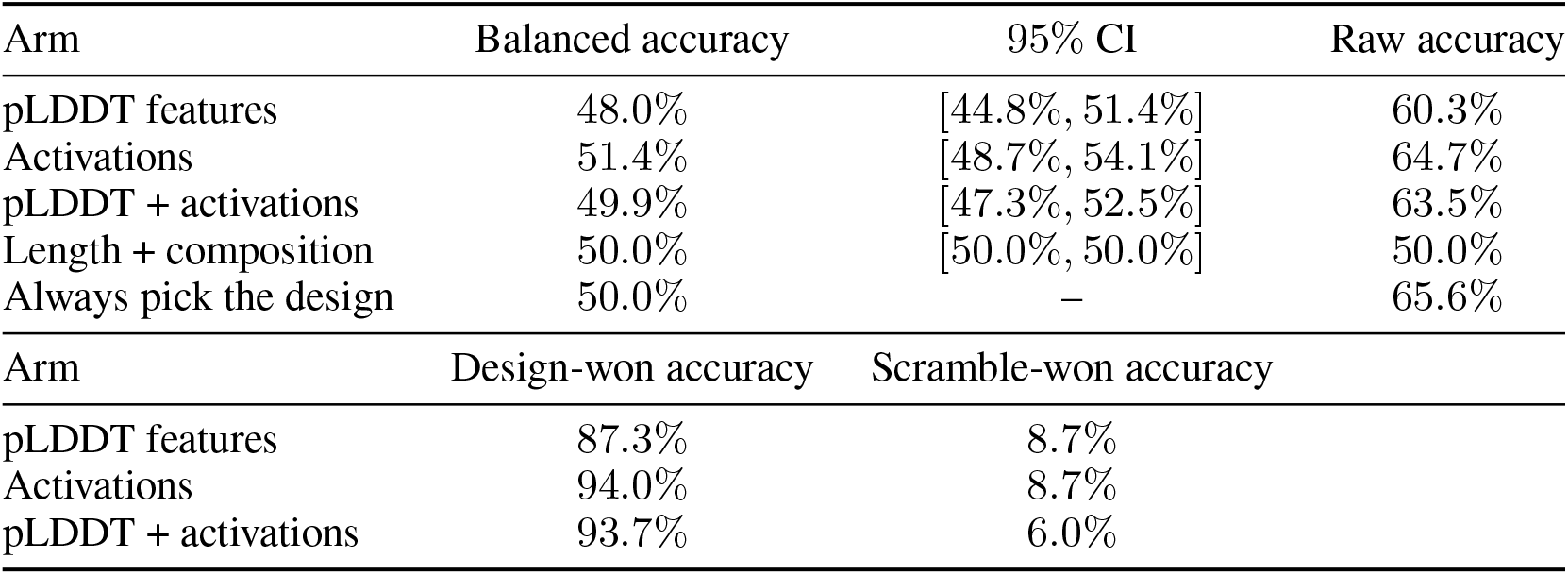
Experimental stability evaluation on *N* = 433 matched design–scramble pairs from the experimental measurements of [13], using nested cross-validation. Pairs match length and composition and differ in amino-acid order. The upper block reports balanced accuracy with 95% confidence intervals and raw accuracy. The lower block separates the 284 design-won and 149 scramble-won pairs. Identical paired composition features force the composition-only arm to tie. No balanced-accuracy interval excludes chance, and no learned arm exceeds the always-design rule’s raw accuracy. The dash denotes an interval not supplied for that rule.

## G Related work

This appendix expands on the related work in Section 1. Our contribution is an audit rather than a new method.

### G.1 Steering protein language models

The method we audit [20] controls repetition by contrastive activation steering, reports its main results at a single operating point, and scores utility as an average of AlphaFold pLDDT and pTM. Its own ablation acknowledges that repetition control saturates while utility declines beyond approximately *α* = 1–2. We add an independent structural readout, a fitted dose threshold with a bootstrap interval and its own control, and matched-norm direction controls at two doses. Contrastive activation steering for proteins is established more broadly [6].

Independent protein work reaches a compatible conclusion by a different route [3]. Comparing their method against two activation-steering baselines, they report substantial quality degradation and a symmetric effect under both addition and ablation, which they read as “off-manifold disruption rather than selective concept control.” Our saturation result—random, coordinate-shuffled and orthogonal directions all indistinguishable from the constructed one at 2 ×—is the same phenomenon measured on repetition rather than toxicity.

### G.2 Direction and identifiability

Prior work establishes that steering directions are not unique in language models [17] by perturbing a vector and comparing *v* with *v* + Δ*v*. We replace the vector entirely with a random direction at matched norm. This complementary test asks whether *any* direction of that magnitude produces the effect, not whether a nearby one does. Direction is irrelevant at saturation but decisive at moderate push in our tests, so neither result generalises to the other dose.

### G.3 The confidence readout

Prior work demonstrated pLDDT’s length sensitivity on AlphaFold in 2022 [11], both as an observed trend and in a controlled random-sequence sweep across lengths. We replicate it on ESMFold using paired within-sequence truncation. Our contribution concerns its consequence for evaluation: if an intervention changes generation length, the readout partly scores that change, which can manufacture a false null.

Prior work also predicts fold confidence from language-model embeddings at far greater scale [2], regressing AlphaFold2 pLDDT over more than a million natural sequences. We study generated rather than curated sequences, use a different structure predictor, and frame prediction as binary triage with abstention rather than regression. We do not improve on that work. These differences make the reported accuracies incomparable, so we do not present them side by side.

The design–scramble evaluation uses measured protease resistance [13], whose stability score already subtracts an unfolded-state prediction fitted on scrambled sequences. That is why comparing a design against its own scramble is not circular.

### G.4 Sparse features and interpretability

Sparse autoencoders are an active line in protein language models [15; 9; 19; 10; 14], spanning feature discovery, semantically guided dictionaries, steering through sparse representations, pair representations in co-folding, and geometric annotation of latent features. This work provides context rather than evidence for our prediction arm.

Prior work shows that sparse features can underperform raw activations across several language-model probing regimes [7]. Our matched-budget comparison replicates this result in proteins, but carries no uncertainty estimate and establishes no significance. The gap narrowed as the dictionary improved, making it an upper bound on the disadvantage rather than a floor.

### G.5 What is ours

Our audit tests a published preservation claim with an independent structural readout, fits a dose threshold with a bootstrap interval and its own control, and compares matched-norm directions at two doses, reporting both outcomes. We use paired within-sequence truncation to examine length’s consequence for intervention evaluation. We also measure triage costs, reporting each saving with its recall, and check both readouts against experimental stability in composition- and length-matched pairs.

## Footnotes

2 Nested cross-validation selected layer 18 (15 of 25 folds), not Section 5’s layer 30.

